# Stable encoding of visual cues in the mouse retrosplenial cortex

**DOI:** 10.1101/785139

**Authors:** Anna Powell, William M. Connelly, Asta Vasalauskaite, Andrew Nelson, Seralynne D. Vann, John Aggleton, Frank Sengpiel, Adam Ranson

## Abstract

The rodent retrosplenial cortex functions as an integrative hub for sensory and motor signals, serving roles in both navigation and memory. While retrosplenial cortex (RSC) is reciprocally connected with the sensory cortex, the form in which sensory information is represented in the retrosplenial cortex and how it interacts with behavioural state is unclear. Here, we used 2-photon cellular imaging of neural activity of putative excitatory (CaMKII expressing) and inhibitory (parvalbumin expressing) neurons to measure visual and running evoked activity in RSC and compare it to primary visual cortex (V1). We found that stimulus position and orientation information was preserved between V1 and RSC, and additionally that positional information was organised topographically. Stimulus directional preference was biased towards nasal-temporal flow. Locomotion modulation of activity of single neurons, both in darkness and light, was also more pronounced in RSC than V1, and strongest in parvalbumin-positive neurons. Longitudinal measurements of single neurons showed that these response features were stably maintained over many weeks. These data provide evidence for stable representations of visual cues in retrosplenial cortex which are highly spatially selective. These may provide sensory data to contribute to the formation of memories of spatial information.

---

The rodent retrosplenial cortex (RSC), which comprises much of the medial part of the dorsal cortex, is reciprocally connected to the hippocampal formation, anterior thalamic nuclei and visual cortex. It is, therefore, perhaps unsurprising that this cortical region has been repeatedly shown to be important for spatial memory and navigation (Vann et al. 2009). However, there is still much uncertainty with respect to the nature of the RSC’s contribution to these processes. One proposal that has been at the forefront of RSC research for the last decade is that this structure has an important role in integrating sensory and non-sensory information (Wolbers and Büchel 2005; Byrne et al. 2007; Vann et al. 2009; Alexander and Nitz 2015); this proposal reflects its anatomical connectivity making it well-placed to combine cortical and subcortical sensory and motor related signals. Consistent with a role in integrating sensory signals, experiments in anesthetised mice provide evidence of sensory responses in the RSC (Murakami et al. 2015; Zhuang et al. 2017) while other work has described visually evoked responses in feedback axons relaying signals between retrosplenial cortex and visual cortex (Makino and Komiyama 2015). Meanwhile, unit recordings in freely moving rats have showing that retrosplenial unit ensembles simultaneously map position in both external and internal frames of reference (Alexander and Nitz 2015).

This integrative model of retrosplenial function is further supported by behavioural data, as RSC lesioned rats are particularly impaired on tasks where they are explicitly called upon to integrate different types of information (Cooper and Mizumori, 1999; Vann and Aggleton, 2002; Pothuizen et al., 2008; Elduayen and Save, 2014; Hindley et al., 2014). Despite the apparent importance of the RSC for spatial cognition, there is surprisingly little evidence from awake animals examining how sensory information required for these functions is encoded in RSC. In particular, the extent to which simple visual stimuli are encoded in an abstract or higher-order vs veridical representation (i.e. which encodes the physical properties of the stimulus) remains unclear. More broadly, it remains uncertain how visual drive is integrated with motor input in RSC. Finally, the extent to which RSC visual and non-visual representations, in individual neurons, are longitudinally stable (as in early sensory cortex) or alternatively dynamic and context sensitive, is unknown.

To address this gap in our knowledge, we used cellular resolution 2-photon imaging, together with genetic labelling of cell types, to assess how simple retinotopic visual and motor related signals are organised and processed in RSC. A key step was to study head-fixed awake animals, allowing precise control and disentanglement of visually driven and motor driven neural activity. This allowed an examination of whether visual and motor signals are processed by overlapping or distinct populations of RSC neurons, and how stable or dynamic representations of these signals are in single neurons over timescales of weeks.

## Materials and Methods

### Animals

All experimental procedures were carried out in accordance with the UK Animals (Scientific Procedures) Act 1986 and European Commission directive 2010/63/EU. Mice expressing GCaMP6F were generated by crossing the Ai95D line (Jax, 024105) with either the CaMkII-alpha-cre T29-1 line (Jax, 005359), for expression in CaMKII positive cells, or the PV-cre line (Jax, 008069), for expression in PV positive cells. Experiments were carried out on adult male mice, housed under normal light conditions (12h light, 12h dark). All recordings were made during the light period.

### Animal surgical preparation

All surgical procedures were conducted under aseptic conditions. Prior to cranial window surgery, animals were administered with the antibiotic Baytril (5mg/kg, s.c.) and the anti-inflammatory drugs Rimadyl (5mg/kg, s.c.) and Dexamethasone (0.15mg/Kg, i.m.). Anesthesia was induced at a concentration of 4% Isoflurane in oxygen, and maintained at 1.5-2% for the duration of the surgery. Once anesthetised, animals were secured in a stereotaxic frame (David Kopf Instruments, Tujunga, CA, USA) and the scalp and periosteum were removed from the dorsal surface of the skull. For RSC recordings, a custom head plate was attached to the cranium using dental cement (Super Bond, C&B), with an aperture approximately centred over the right hemisphere retrosplenial cortex. A 3mm circular craniotomy was then made, centred approximately 2.5mm caudal to bregma. The lateral placement of the craniotomy was such that it encompassed a small portion of the left hemisphere. This positioning meant that is was possible to visualise the central sinus through the cranial window, which provided a useful reference point when imaging. The craniotomy was closed with a glass insert made from 3 layers of circular glass (#1 thickness; 1×5 mm, 2×3 mm diameter) bonded together with optical adhesive (Norland Products; catalogue no. 7106). The window was placed such that the smaller pieces of glass were in contact with the brain surface and the larger piece rested on the skull surrounding the craniotomy. The window was then sealed with dental cement. After surgery, all animals were allowed at least 1 week to recover before being imaged. For V1 recordings, procedure was performed as described above except the craniotomy and head plate position was centred −3.4 mm posterior and 2.8 mm lateral from bregma of the right hemisphere.

### Imaging and locomotor data acquisition

In vivo 2-photon imaging was performed using a resonant scanning microscope (Thorlabs, B-Scope) with a 16x 0.8NA objective (Nikon). GCaMP6 was excited at 980nm using a Ti:sapphire laser (Coherent, Chameleon) with a maximum laser power at sample of 50mW. Data were acquired at approximately 60Hz and averaged, resulting in a framerate of approximately 10Hz. For all imaging experiments animals were head-fixed and free to run on a custom designed fixed axis cylindrical treadmill. Movement was measured using a rotary encoder (Kübler, 05.2400.1122.0100). Imaging, behavioral and visual stimulation timing data were acquired using custom written DAQ code (Matlab) and a DAQ card (NI PCIe-6323, National Instruments).

### Visual stimuli

Visual stimuli were generated in Matlab using the psychophysics toolbox (32), and displayed on 2 calibrated LCD screens (Iiyama, B2080HS; width x height 26 x 47 cm) at right angles to one another () placed 20 cm from the eye. All visual stimuli were circular, unidirectionally drifting square wave gratings of uniform size (40×40 deg), spatial (0.08 cycles per degrees) and temporal frequency (1 Hz) presented at full contrast. Stimulus position was corrected for viewing angle. Stimulus parameters were optimised in pilot experiments. Orientation and spatial location varied depending on the experiment. For all experiments, drifting gratings were presented for two seconds followed by a two second inter-trial interval during which a gray screen was displayed. In all experiments visual stimuli were presented passively, such that the drifting of the grating was not yoked to the animal’s movement.

### Experimental design

Low magnification imaging of the entire extent of RSC visible within the cranial window was used to localise the visually responsive region. In order to elicit visually driven activity, a vertically orientated drifting grating, was presented in the centre of the binocular region of the visual field, centred at 20 deg elevation. Recordings were made at 11 different depths from pial surface to −300µm, at two partially overlapping 900×900µm fields of views within the RSC (one rostral and one caudal).

For experiments to determine spatial and orientation selectivity, 48 different stimuli presented, with each stimulus repeated 10 times in a pseudo-random order. The stimuli were circular drifting gratings of one of eight different orientations, presented in one of six locations in a 3 x 2 grid, centered at 0, 45 and 90 degrees in azimuth, and 0 and 20 degrees in elevation. Recordings were made from a 400×400µm field of view, aligned on the medial side with the midline, and centered approximately 1.75mm and 3.25mm caudal of bregma for the rostral and caudal recording sites respectively. Recordings were made at depths of 150-250µm.

### Calcium imaging data analysis

Calcium imaging data was registered and segregated into neuronal regions of interest using Suite2P (Pachitariu et al. 2016). Pixel-wise stimulus preference maps were constructed by first calculating the mean of the registered imaging frames recorded during the drifting phase of each stimulus, and then determining for each pixel the stimulus which elicited the largest mean response.

For experiments in which the stability of visual responses was tracked longitudinally, recordings were made from the same field of view over multiple sessions. Cortical surface vascular landmarks were used to locate the same neurons between sessions. In order to match region of interest masks detected with Suite2P between imaging sessions, the cpselect, fitgeotrans and imwarp functions in Matlab were used to warp the ROI masks using manually identified control points selected from the mean image frame from each experiment. Masks which overlapped by more than 60% and were detected in all sessions were flagged as potential longitudinally recorded neurons. These masks were then manually verified by visual observation of the region of the mean each frame that they corresponded to in each experiment.

### Quantification and statistical analysis of visual and motor responses

The visual responses of individual neurons were quantified as the mean dF/F value between 0.5-1.5 secs after stimulus onset, with a baseline period quantified for each trial as the mean dF/F between 1-0 secs before stimulus onset. The response to drifting gratings over the visual field was fit in MATLAB using a custom model and the fitnlm function. Specifically, the spatial extent of the response was modelled with a 2D Gaussian, and the response to orientation was fit by the sum of two Von Mises functions, separated by 180 degrees. The two Von mises functions were normalized between 0 and 1, and multiplied by the 2D gaussian so that the parameters of the Von Mises function solely reflected orientation selectivity, and the parameters of the 2D Gaussian solely reflected the spatial extent of the receptive field, and the maximum response. Orientation tuning curves were calculated using data pooled over the two directions of grating drift. Orientation tuning fit curves were used to calculate OSI defined as (peak response – trough response)/peak response.

A neuron was classified as visually responsive if it showed a statistically significant increase in activity between the baseline and visual stimulation periods for at least one stimulus, with statistical significance calculated with a shuffle test (1000 shuffles, p<0.05), corrected for multiple comparisons using the mafdr false discovery rate function in Matlab. A one-way ANOVA of response amplitudes over stimulus position, orientation or direction was used to assess whether neurons discriminated these stimulus features. Statistical significance of correlations between run speed and neural activity were calculated with a shuffle test in which the entire trace of run speed was randomly shifted relative to neural activity traces 1000 times (p<0.05). Where appropriate similarity of variance and normality of distribution were checked with the vartestn Matlab function and the Kruskal-Wallis test was used as noted when the assumptions of the one-way ANOVA or t-test were not met. T-tests used were two-sided. Correction of p values for multiple comparisons were calculated using the Matlab function multcompare using the Tukey–Kramer method.

## Results

### Spatial organisation of visually responsive cells in RSC

We first used low magnification 2-photon imaging of GCaMP6f expressed in CaMKII positive cells (i.e. primarily excitatory neurons, Supplementary Figure 1A) or parvalbumin expressing inhibitory neurons (PV neurons, Supplementary Figure 1B) to localise visually responsive areas of RSC. Recordings were made in awake animals free to walk on a cylindrical treadmill (Figure 1A). We sampled the region between −2.4mm and −4.2mm AP and 0.0mm to 0.9mm ML relative to bregma, and depths between the pial surface and 300µm. The area was tiled by two partially overlapping 900×900µm fields of view (one anterior, one posterior), and data were acquired at depths spaced by 30µm in z (Supplementary Figure 1C), resulting in 11 depths per field of view. We first generated mean pixel-wise response maps to binocularly presented drifting grating visual stimulation and collapsed across the 11 depths.

**Figure 1.**
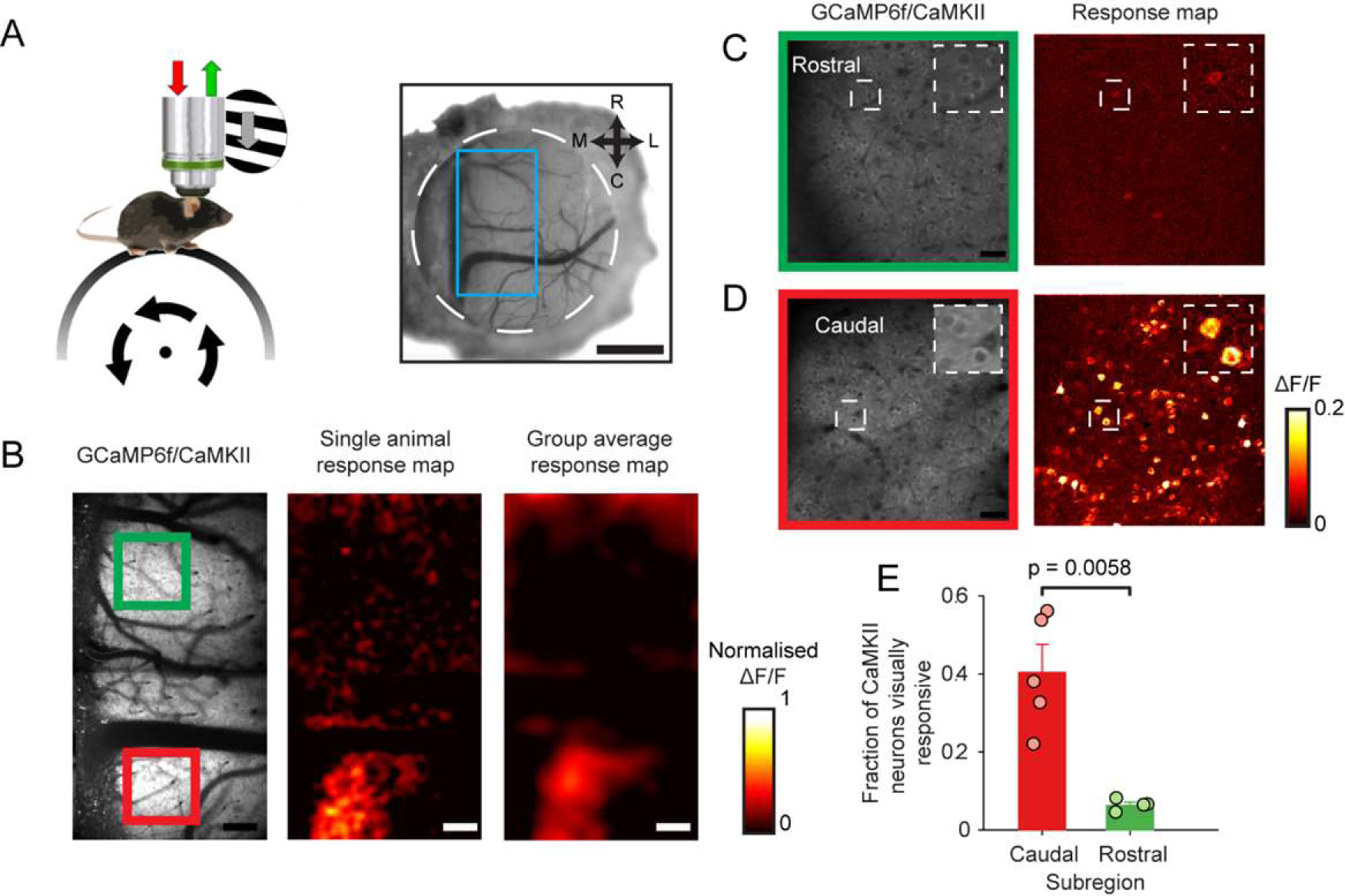
Localization of visually responsive area of dysgranular retrosplenial cortex. (A) Setup schematic (left) and cranial window over retrosplenial cortex (right). Blue box indicates area imaged. Scale bar 1mm. (B) Low magnification 2-photon visual response maps showing visually evoked activity of CaMKII neurons in the caudal (red box) but not rostral (green box) retrosplenial cortex. Scale bar 160µm. (C-D) Higher magnification of rostral (C) and caudal (D) regions showing visually responsive neurons in caudal but not rostral areas. Scale bar 30µm. (E) Mean fraction of visually responsive neurons (error bars represent S.E.M.) detected in caudal and rostral areas.

Response maps of the CaMKII RSC population showed a visually responsive area limited to the caudal RSC (cRSC) that was visible both in maps from individual animals and in the mean response maps averaged across 5 animals (Figure 1B). Analysis of response maps across the 11 depths (spanning 300µm) showed no systematic variation with depth in the visual responsiveness of this caudal region (Supplementary Figure 1D). Response maps of the PV RSC population showed a similar pattern of greater visual responsiveness in the caudal vs. rostral RSC, but with significantly lower amplitude (Supplementary Figure 1E). We next made higher resolution recordings from layer 2/3 of the caudal RSC (cRSC) and rostral RSC (rRSC) in which individual CaMKII neurons could be resolved (Figure 1C and D). Consistent with our low magnification observations, higher magnification pixel-wise maps showed a large fraction of visually responsive cells in cRSC but not rRSC regions (Figure 1E; see also Supplementary Figure 2A). A cell-wise analysis of visual responses of neurons confirmed that a significantly larger fraction of caudal than rostral neurons were visually responsive (fraction of neurons visually responsive: 40.4±6.4% caudal vs. 6.3±7.4% rostral; t-test: p = 0.0058; n = 5 and n = 5 mice respectively). These results show that visually responsive CaMKII and PV neurons are largely confined to caudal regions of the RSC and that within cRSC a larger fraction of neurons is visually responsive than previously reported in anesthetised animals (Murakami et al. 2015).

### Spatial and orientation selectivity of visually responsive cells in cRSC

We next investigated the response properties of individual CaMKII neurons in the visually responsive cRSC. A recent study using widefield imaging suggested the presence of a retinotopic map in the RSC (Zhuang et al. 2017). We therefore first tested for the existence of retinotopy at single cell resolution. Animals were visually stimulated with 40×40 deg grating patches at one of 6 retinotopic locations and 8 orientations. Pixel-wise maps were then calculated for azimuth stimulus preference (Figure 2B). This showed a broad range of retinotopic preferences within the 400×400µm field of view which included the full range of stimulus positions presented. Retinotopic organisation was apparent in the pixel maps of azimuth (which we sampled over a broader range than elevation due to restrictions in possible display positions, Figure 2B and C; mean correlation between azimuth preference and distance from midline = 0.85±0.07; n = 5 mice), and individual neurons typically exhibited spatially selective responses (Figure 2D and Supplementary Figure 2B). In the azimuth map more medial and lateral parts of the map corresponded to more binocular (i.e. nasal) and monocular (i.e. temporal) visual space, respectively. We next compared the spatial selectivity of cRSC CaMKII neurons to that of CaKII primary visual cortex (V1) neurons stimulated with the same stimulus. Of those cRSC and V1 neurons that were visually responsive, a similar fraction discriminated stimulus position (fraction of neurons discriminating stimulus position: 73.7±0.7.1% cRSC vs. 72.4±5.0% V1; t-test: p = 0.997; n = 5 and n = 5 mice respectively; Figure 2E).

**Figure 2.**
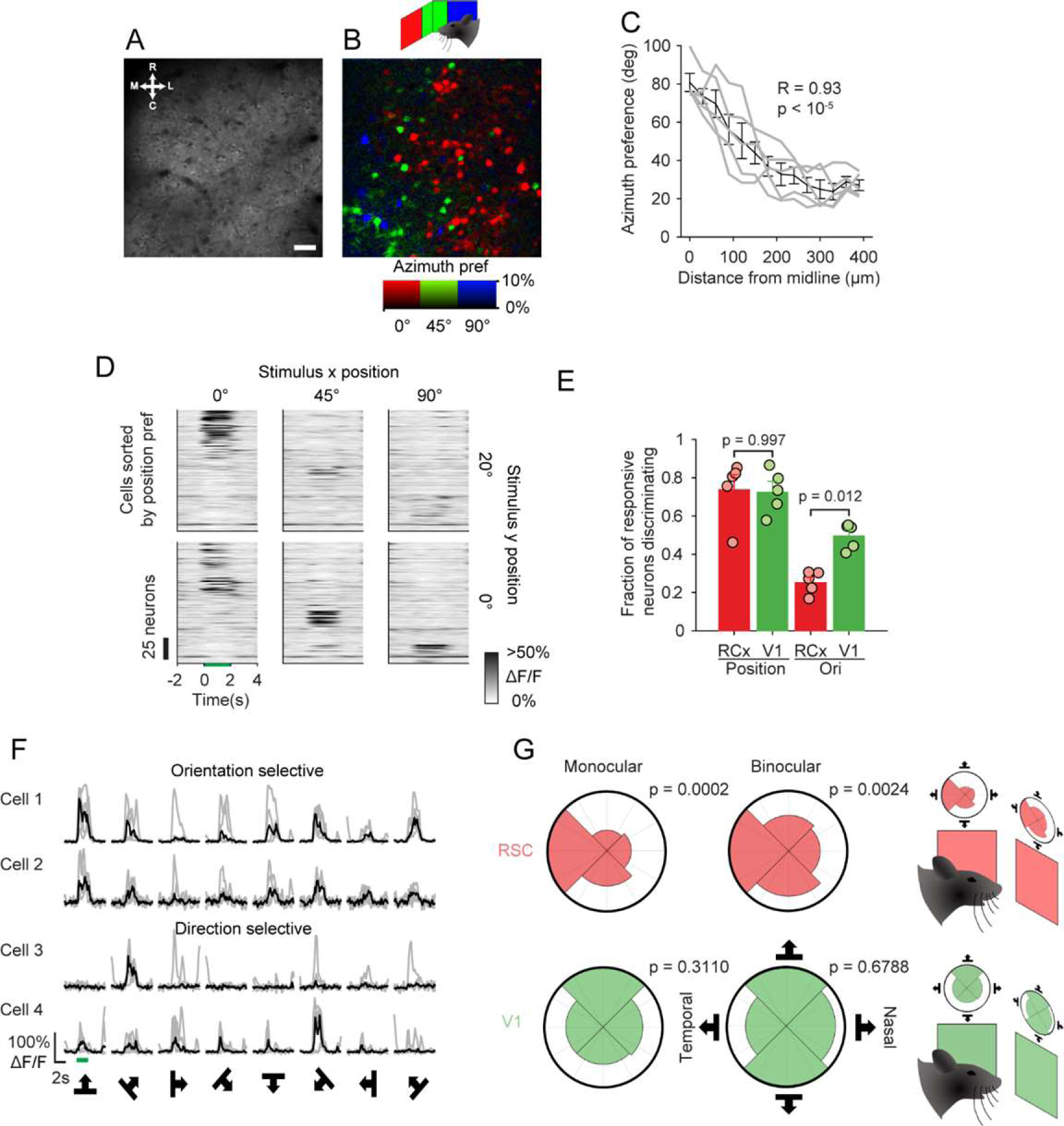
Visual selectivity of CaMKII cRSC neurons. (A-B) Mean imaging frame from cRSC (A), and pixel-wise map from the same field of view of retinotopic preference in azimuth (B). Scale bar 30µm. (C) Azimuth position preference compared to distance of neurons from midline. Grey lines represent individual animals and black line is average. (D) Mean raster plot of neural responses to stimuli at different retinotopic locations, with neurons (rows) sorted by positional preference. (E) Mean fractions of neurons (error bars indicate S.E.M.) statistically significantly discriminating stimulus position or orientation in cRSC and V1. (F) Example traces of orientation selective and direction selective neurons. (G) Polar plots showing summary of directional preference of all neurons recorded in RSC (red) and V1 (green) divided into those representing the monocular and binocular visual field. P values indicate statistical significance of Raleigh test of non-uniformity and show non-uniformity of RSC direction preference distribution.

In contrast we found that only 25.2±2.6% of visually responsive cRSC neurons significantly discriminated stimulus orientation or direction (at preferred position; Figure 2F) which compared to 49.5±0.03% of V1 neurons (t-test: p = 0.012; n = 5 and n = 5 mice respectively; Figure 2E). This was reflected in a median orientation selectivity index of 0.63 in cRSC as compared to 0.81 in V1 (Kruskal–Wallis test: p < 10^-12^, n = 436 and n = 307 neurons respectively). We additionally observed a significant bias in directional preference of orientation selective neurons cRSC neurons (defined as neurons with OSI > 0.5) towards naso-temporal motion (Figure 2G). This bias was not observed in orientation selective V1 neurons, and was apparent both in neurons encoding monocular visual space (Raleigh test for non-uniformity; V1: n = 70 neurons; p = 0.3110; RSC, n = 115 neurons, p = 0.00017) and binocular visual space (V1: n = 219 neurons, p = 0.68; RSC: n = 202 neurons, p = 0.0024), although it appeared more pronounced in monocular neurons. Finally, we observed that the degree of habituation of responses over the course of the experiment of visually responsive neurons to their preferred stimuli differed between V1 and RSC neurons, with RSC neurons exhibiting a greater reduction in ΔF/F than V1 neurons (reduction of 0.35 vs. 0.11; t-test: p = 0.001; Supplementary Figure 2D). Together these data show that a significant fraction of cRSC neurons show response selectivity for retinotopy, orientation and direction of visual flow and as a whole the population is organised retinotopically.

### Integration of visual cues and locomotion signals in RSC

Sensory processing by V1 neurons is modulated by the locomotor state of the animal (Niell and Stryker 2010; Keller et al. 2012; Saleem et al. 2013; Ranson 2017; Dipoppa et al. 2018).

We next examined the extent to which RSC sensory processing is subject to the same modulation, and the way in which motor signals are integrated with sensory signals. We first performed recordings of RSC and V1 CaMKII neurons in complete darkness to isolate locomotor from visual stimulus related activity. We found that, in both RSC and V1, a fraction of neurons was significantly correlated with run speed (either positively or negatively; shuffle test, p < 0.05; Figure 3A-C). Both caudal and rostral RSC exhibited a similar pattern of run speed correlations in darkness and so were combined in this analysis (see Supplementary Figure 3A and B for data split into rostral and caudal regions). A larger fraction of RSC than V1 CaMKII neurons was found to be significantly positively correlated with run speed (25.9±0.02% RSC vs. 10.8±0.02% V1; t-test: p = 0.0003; n = 10 fields of view [5 mice] and n = 6 fields of view [3 mice] respectively, Figure 3D and E) or significantly negatively correlated with run speed (17.1±0.02% RSC vs. 10.5±0.02% V1; t-test: p = 0.0338; n = 10 fields of view [5 mice] and n = 6 fields of view [3 mice], Figure 3D and E). Consequently, some RSC neurons were found to be active in darkness and reliably suppressed by locomotion (Figure 3A upper rows), while other neighbouring neurons recorded simultaneously exhibited the opposite behaviour (Figure 3A lower rows). We next recorded from PV RSC neurons under the same dark conditions (Figure 4A). A similar distribution was again observed in caudal and rostral RSC and so neurons from the two areas were again pooled (see Supplementary Figure 4A and B for data split into rostral and caudal regions). In contrast to the CaMKII population (Figure 4B), PV neurons exhibited an overall higher degree of locomotion correlation (Figure 4C), both in neurons which were positively correlated (median R = 0.19 CaMKII vs. 0.34 PV; Kruskal–Wallis test p < 10^-8^; Figure 4B, C and D) and in negatively correlated neurons (median R = −0.13 CaMKII vs. −0.26 PV; Kruskal–Wallis test: p < 10^-14^; Figure 4B, C and E). The distribution of correlation coefficients of significantly negatively correlated neurons was found to be bimodal suggesting functional diversity in the PV population (Figure 4C).

**Figure 3.**
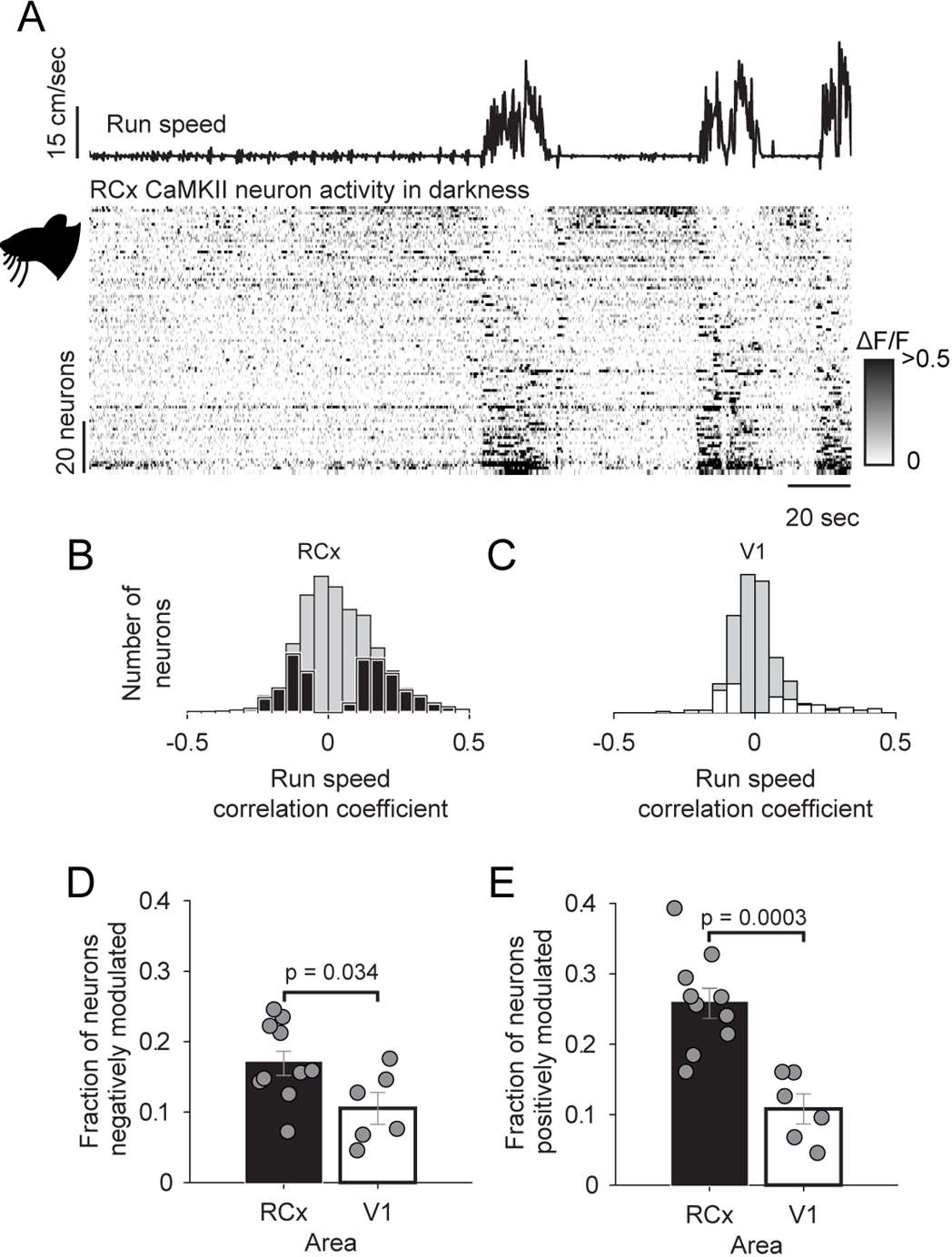
Comparison of the activity of CaMKII RSC and V1 neurons in darkness. (A) Run speed, and below, raster representation of neural activity (rows are individual neurons) sorted by run speed correlation. (B-C) Histogram of run speed correlations of RSC neurons (B, black) and V1 neurons (C, white) with the non-significantly correlated fraction of neurons (determined with shuffle test) represented by gray bars. (D-E) Mean fraction of neurons which are significantly negatively locomotion correlated (D) or positively locomotion correlated (E). Error bars indicate S.E.M.

**Figure 4.**
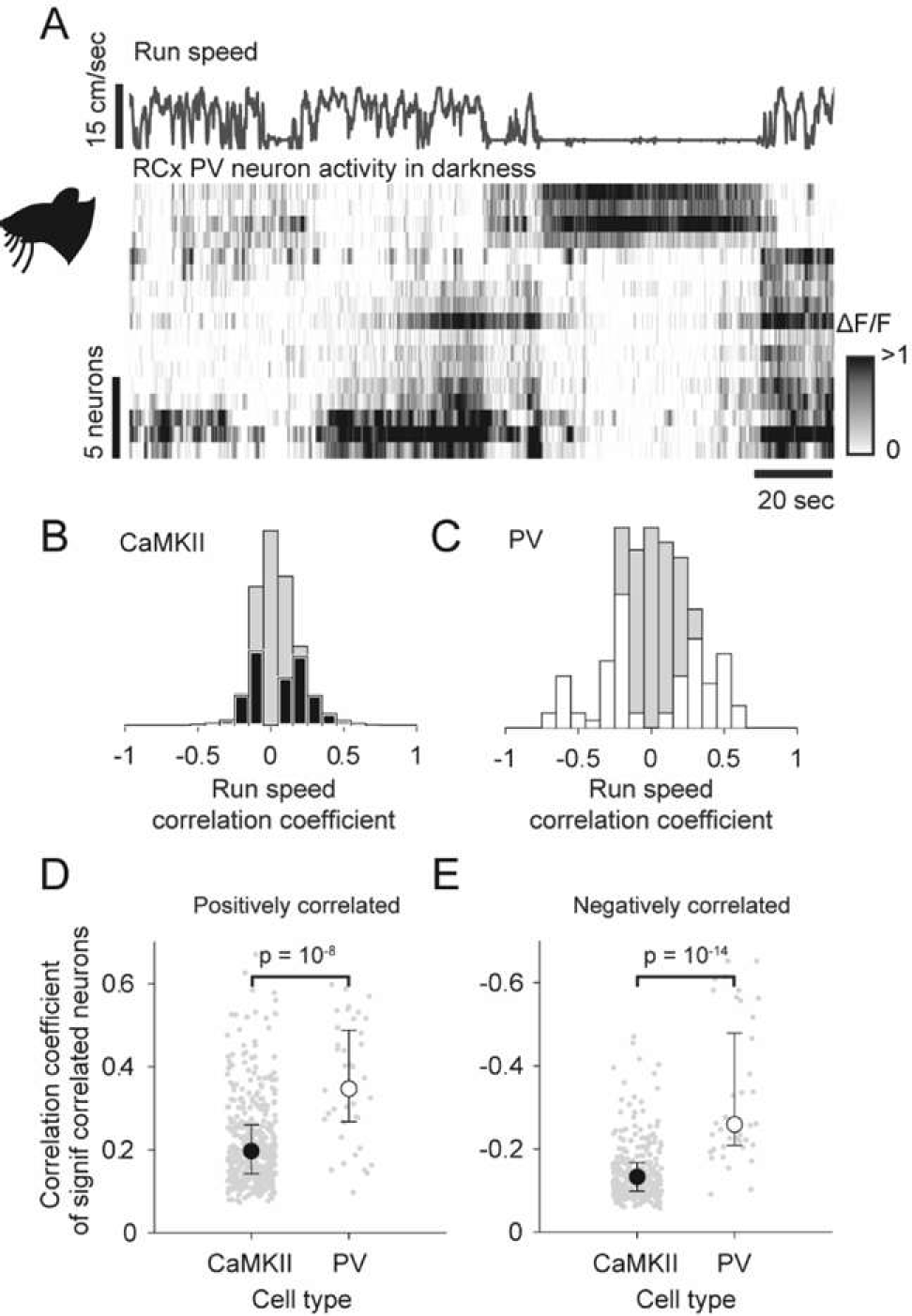
Activity of PV RSC neurons in darkness. (A) Run speed, and below, raster representation of neural activity (rows are individual neurons) sorted by run speed correlation. (B-C) Histogram of run speed correlations of CaMKII neurons (B, black) and PV neurons (C, white) with the non-significantly correlated fraction of neurons represented by gray bars. (D-E) Median fraction of neurons which is significantly positively locomotion correlated (D) or negatively locomotion correlated (E). Error bars indicate interquartile range.

We next sought to determine the interaction between visual input and locomotion related input in individual cRSC neurons (Figure 5A-D). We first quantified the extent to which the visually responsive cRSC population overlaps with the locomotion correlated population. To do this we compared the run speed correlation coefficients of visually responsive vs. non-visually responsive cells. We found no significant differences between these groups, either in CaMKII neurons (Supplementary Figure 5A) or PV neurons (Supplementary Figure 5B), suggesting no correlation between visual stimulus sensitivity and locomotion sensitivity in individual cRSC neurons. As was the case in darkness, we found that PV neurons on average exhibited a higher degree of correlation with run speed than CaMKII neurons during visual stimulation, both in neurons that were positively correlated (median R = 0.11 CaMKII vs. 0.16 PV; Kruskal–Wallis test: p < 10^-4^; Figure 5E left panel) and in negatively correlated neurons (median R = −0.06 CaMKII vs. −0.15 PV; Kruskal–Wallis test: p < 10^-12^; Figure 5E right panel). We next investigated the effect of locomotion on the responses of cRSC neurons to visual stimuli. To quantify this we calculated an index of locomotion modulation (LMI) defined as: (moving response – stationary response)/(moving response + stationary response), calculated with responses to stimuli of preferred orientation and position. This index ranges in value between −1 and 1, and larger magnitude values indicate a greater modulation. The majority of cRSC CaMKII neurons had a positive LMI (81%), the median LMI was 0.40 (Figure 5F), and the distribution was unimodal. In contrast, the distribution of LMIs of cRSC PV neurons was bimodal with a significant fraction of cells exhibiting either strong suppression or strong facilitation of visually evoked activity by locomotion (Figure 5G), again suggesting a significant functional diversity in the cRSC PV population. In summary, in darkness RSC neurons are significantly more locomotion modulated than V1 neurons, and within the RSC population PV neurons are more locomotion modulated than CaMKII neurons, with a subset of PV neurons particularly strongly suppressed during locomotion. Both visual and on-visual RSC neurons are similarly influenced by locomotion, and the degree of locomotion modulation of visual responses in CaMKII neurons is comparable to that observed in V1.

**Figure 5.**
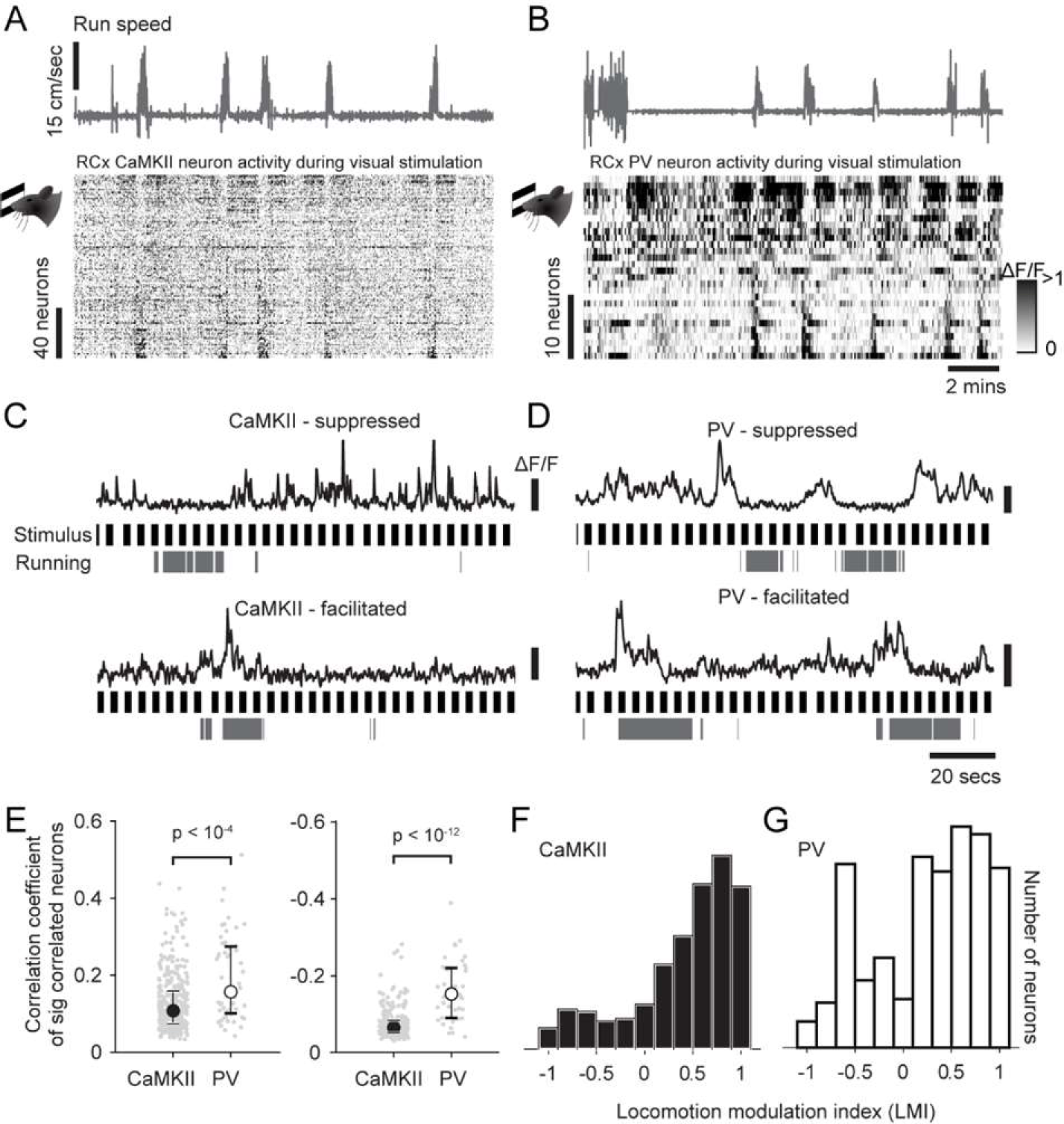
Interaction of visual and locomotion induced activity in CaMKII and PV neurons. (A-B) Run speed, and below, raster representation of neural activity during visual stimulation sorted by run speed correlation in CaMKII neurons (A) and PV neurons (B). (C-D) Example traces of running suppressed (upper) and running facilitated neurons (lower) during visual stimulation, together with running periods and stimulus onset times (black) in CaMKII neurons (C) and PV neurons (D). (E) Median run speed correlations during visual stimulation of significantly positively and negatively correlated neurons (left and right respectively). Error bars indicate interquartile range. (F-G) Histogram of indices of locomotion modulation of visually evoked activity of CaMKII (black, F) or PV (white, G) neurons.

### Long-term stability of RSC visual and locomotion signals

The degree of long term stability of sensory evoked activity varies between brain regions (Clopath et al. 2017), with early sensory areas tending to be on the whole more stable (Mank et al. 2008; Margolis et al. 2012; Rose et al. 2016; Ranson 2017; Jeon et al. 2018) while regions containing more abstract representations, such as regions of the hippocampus, exhibit more dynamic and context sensitive representations (Mankin et al. 2012; Ziv et al. 2013). As the RSC occupies an intermediate position between early sensory cortex and the hippocampus, we next examined the extent to which sensory evoked activity and locomotion modulation of RSC activity was stable over timescales of weeks.

To measure stability of visually evoked activity we focussed on cRSC and the CaMKII population as this was the area and cell type in which we observed visually evoked activity. Over the course of 4 weeks we made 3 evenly spaced recordings of the same CaMKII cRSC neurons, relocated from session to session using vascular landmarks. Pixel-wise maps of stimulus preference showed clear similarity of retinotopic preference in all animals measured over the 4 week period (Figure 6A). To quantitatively assess similarity, we segregated the field of view of each experiment into neuronal regions of interests (ROIs) and validated each ROI as present and valid in each session. This resulted in 170 unambiguously identified CaMKII neurons from 3 animals that were visible in all 3 sessions (mean neurons per animal 56.7±9.9, see Supplementary Figure 6A). We next measured the fraction of neurons with significant visual responses over the 3 sessions and found that the largest fraction (36.5%) had significant responses in all 3 sessions with 19.4% responsive in 2 sessions, 27.7% responsive in 1 session, and 16.5% of neurons not responding in any session (Figure 6B). For each neuron we then calculated the difference between its fitted azimuth retinotopic preference for each pair of sessions and, consistent with the observed similarity of pixel maps, found that the majority of neurons (65%) had differences in positional preference of less than 10 deg (mean difference 7.67±0.86, compared to mean difference with shuffled neuron identities 34.97±0.38; Figure 6C). We next compared the similarity of orientation preference in longitudinally identified orientation selective neurons (OSI>0.5) over all pairs of sessions and found that while the largest fraction of neurons (42.3%) had orientation preferences which differed by less than 10 degrees between sessions, many neurons had large differences in preference (median difference 15.56 degrees, Figure 6D). This difference in tuning preference is considerably larger than previous reports from awake mouse V1 of median differences in orientation preference of approximately 3-4 degrees over a one week period (Jeon et al. 2018). In summary, these results show longitudinal stability of visual response tuning over periods of weeks in many cRSC neurons, particularly with respect to retinotopic preference, and to a lesser degree orientation preference.

**Figure 6.**
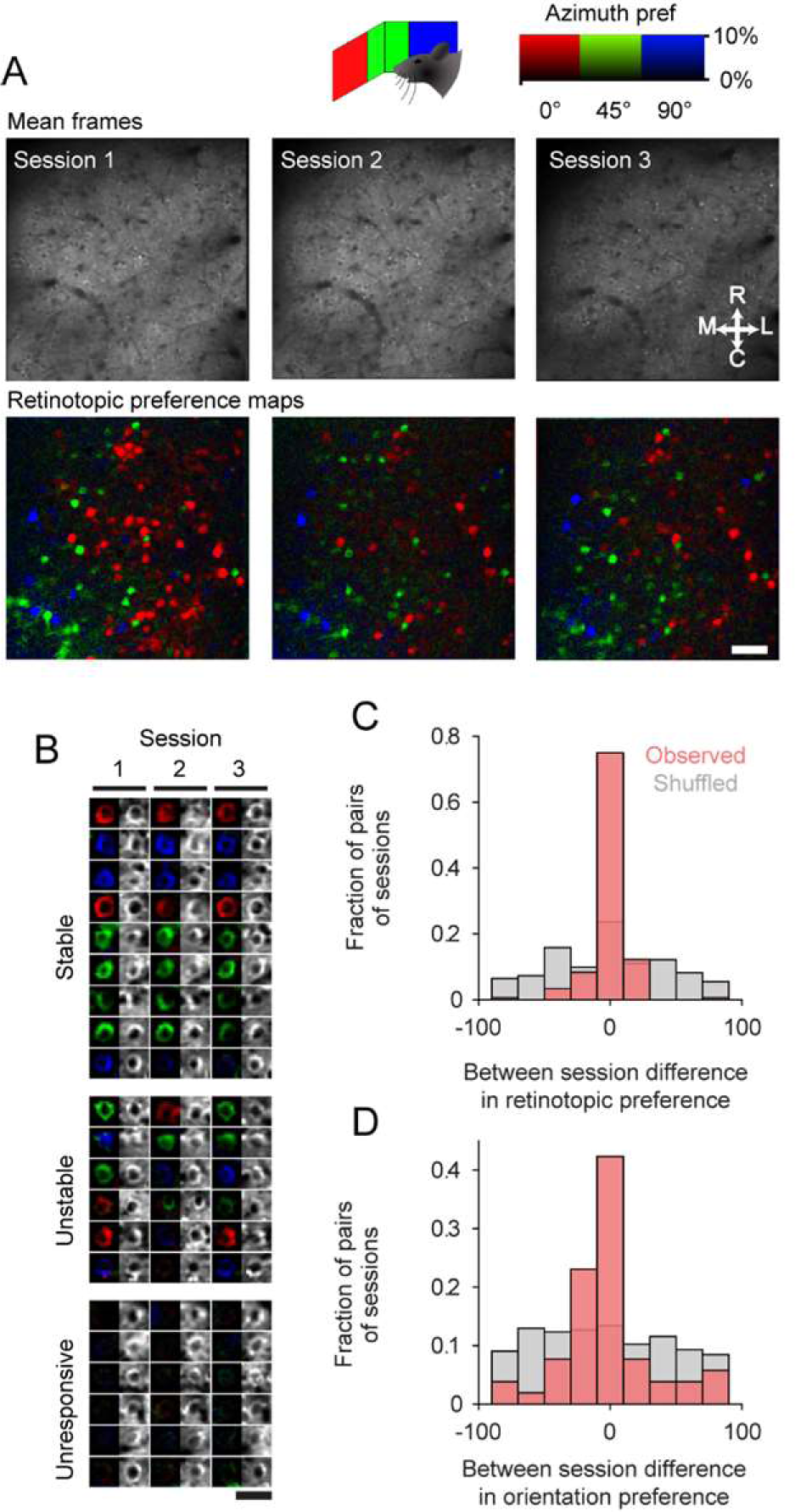
Longitudinal stability of retinotopic positional preference in cRSC neurons over weeks. (A) Single field of view imaged over 3 sessions spread over 4 weeks, showing mean imaging frame (above) and pixel-wise azimuth retinotopic preference map (below). Scale bar 30µm. (B) Examples of longitudinally tracked neurons which has stable visual responses, unstable responses or were stably unresponsive. (C-D) Between session difference in retinotopic preference (C) and orientation preferences (D) for each neuron on each pair of sessions on which it was measured.

We next sought to determine whether the degree of locomotion modulation of individual neurons was also stable over time. As both CaMKII and PV cRSC neurons exhibited clear locomotion modulation we analysed both populations with a spacing of 1-2 weeks between recordings. We used the same criteria outlined above to identify PV neurons which were unambiguously detected in all recording sessions, and this resulted in 52 longitudinally tracked cells from 3 animals (mean neurons per animal 17.33±1.8; see Supplementary Figure 7-B) as well as the 170 CaMKII neurons described above (Supplementary Figure 7-A). We generated pixel-wise maps of locomotion modulation by calculating for each pixel (moving frames – stationary frames)/(moving frames + stationary frames) and then segregated this map into the ROIs which could be identified longitudinally (Figure 7A; Supplementary Figure 7-A). Neurons within cRSC of both cell types exhibited a range of degrees of locomotion modulation, and a high degree of inter-session stability could be seen in many neurons (Figure 7A, columns 1 and 2). We next segregated the field of view into longitudinally tracked neurons and calculated run speed correlations of each neuron in the 2 sessions (Figure 7B and C). This also revealed a striking degree of stability of run speed correlation between sessions that was most pronounced in PV neurons (R = 0.78; Pearson’s correlation coefficient: p < 10^-11^), but also highly significant in CaMKII neurons (Pearson’s correlation coefficient: R = 0.41, p < 10^-7^).

**Figure 7.**
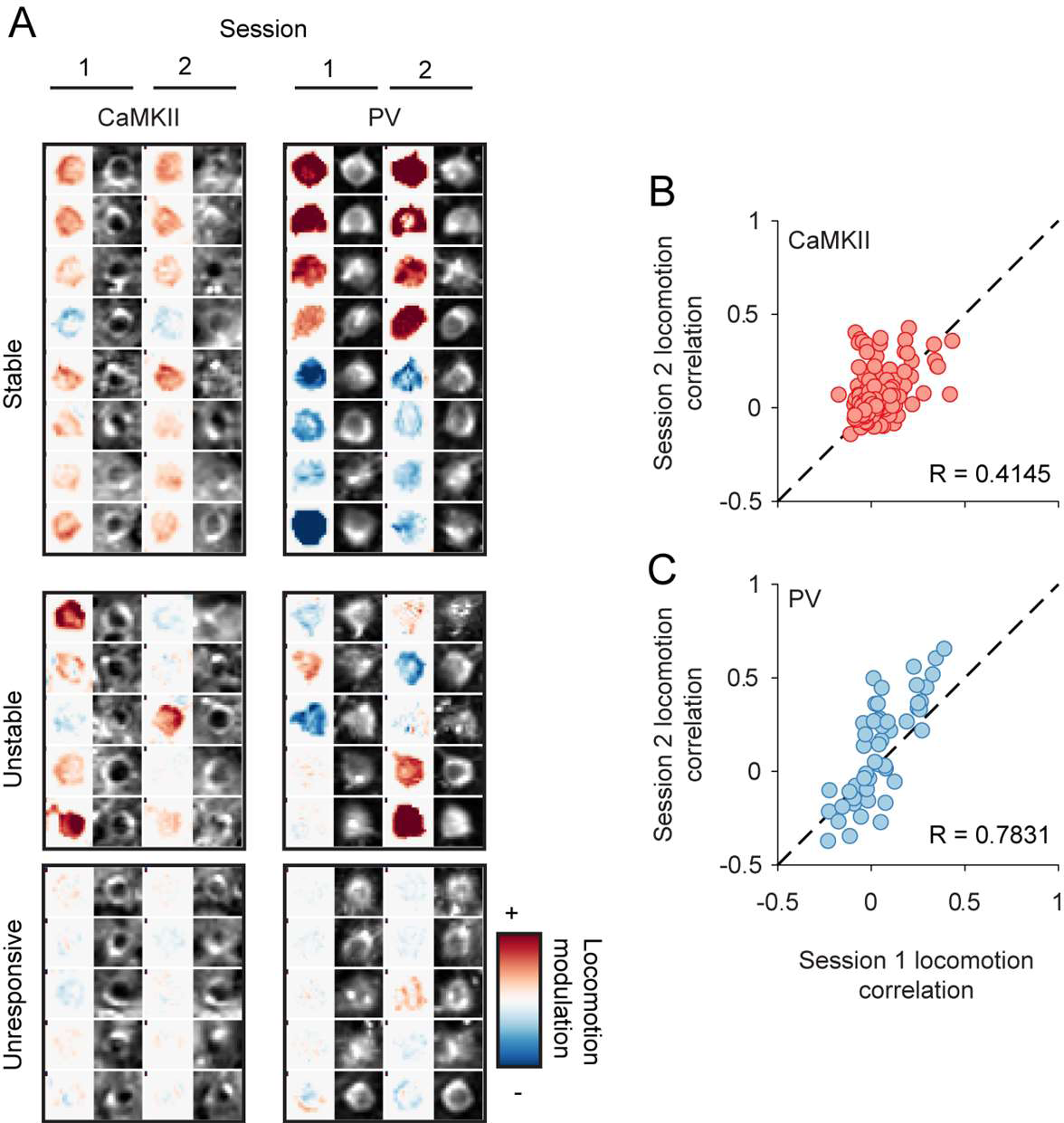
Longitudinal stability of locomotion correlation of cRSC neurons over weeks. (A) Examples of longitudinally tracked CaMKII neurons (left) or PC neurons (right) which had stable locomotion modulation, unstable modulation or were stably unmodulated. (B-C) Inter-session comparison of locomotion modulation of CaMKII neurons (B) or PV neurons (C).

In summary, these results show that a large fraction of RSC neurons exhibit a high degree of stability of both sensory responses and modulation by locomotion. PV neurons were observed to exhibit a particularly high correlation with locomotion and also a high degree of stability of run speed correlation over the timescale measured.

## Discussion

The RSC has been suggested to function as an integrative hub for sensory and motor signals. Here we examined the organisation of visually evoked activity within the RSC in awake mice, how it relates to locomotion related activity, and how stable these motor and visual representations are over periods of weeks.

We observed that a substantial population of dysgranular RSC excitatory neurons exhibit visually evoked activity, and that this subpopulation is largely limited to the more caudal RSC. The dysgranular RSC has gradients of connections that are consistent with this observation. In particular, caudal dysgranular RSC is more interconnected (both afferent and efferent) with primary and higher cortical visual areas (Wyss and Van Groen 1992). Consequently, these results suggest that visual responsiveness is concentrated in those RSC areas with direct visual inputs, i.e., the intrinsic connections linking different parts of dysgranular RSC (Jones et al. 2005) are not sufficient to enable visual responsiveness as measured in the present study. Likewise, it is the caudal dysgranular RSC that shows the clearest difference in *c-fos* activity when contrasting spatial behaviours in the light and the dark (Pothuizen et al. 2009).

We further found that many visually responsive cRSC CaMKII neurons exhibited retinotopic, orientation and direction selectivity. This was largely stably preserved over periods of several weeks. The observation of a high degree of retinotopic selectivity is perhaps surprising given previous evidence of the importance of dysgranular RSC in tasks dependent on allocentric encoding (Vann and Aggleton 2005) as well as recent evidence of place field like activity (Mao et al. 2017). Our observation of cellular resolution retinotopic organisation of cRSC is consistent the view that RSC contains sensory representations of an egocentric nature, by which it is meant that stimuli in specific locations defined by their position relative to the observer can activate specific RSC cells. Therefore, in future work it will be important to reconcile how the retinotopically structured map of visual space reported here, interacts with any equivalent allocentric maps of space. The bias in directional preference of cRSC neurons towards naso-temporal motion may indicate a role in the role in the processing of visual flow information. Computational models show that visual flow provides information about egocentric motion and influences firing patterns in spatially tuned cells during rodent navigation (Raudies et al. 2012). Head direction cells are present in the rodent retrosplenial cortex (Chen et al. 1994), therefore this area may play a role in updating head orientation during movement.

A large number of studies in mice have reported locomotion modulation of both baseline and sensory evoked cortical activity in early sensory areas, including the primary visual and auditory cortices, with the direction of effects varying between cortical areas, stimulation conditions, and cell types (Niell and Stryker 2010; Ayaz et al. 2013; Polack et al. 2013; Saleem et al. 2013; Benevento et al. 2016; Dipoppa et al. 2018; Shimaoka et al. 2018). Our recordings of RSC in the dark revealed that a significant fraction of dysgranular RSC neurons also exhibit baseline activity, which is either strongly positively or negatively correlated with locomotion. As rRSC is more strongly connected than cRSC to both primary motor and secondary motor cortex (Wyss and Van Groen 1992; Shibata and Naito 2008; Yamawaki et al. 2016), we might have expected to see the strongest correlates of locomotion in rostral regions of RSC. Instead, we observed comparable locomotion related activity across the rostral and caudal RSC subregions, which may indicate that locomotion correlations are driven by other motor or arousal related signals such as neuromodulatory input (Fu et al. 2014). These findings may also relate to previous studies showing the importance of the RSC for path integration which relies on locomotion-derived cues to update internal representations of space (Cooper and Mizumori 1999, 2001; Elduayen and Save 2014).

Interestingly, the visual responsiveness and locomotion modulation of cRSC neurons were found to be largely unrelated to one another, with both highly visual and visually non-responsive neurons exhibiting similar levels of locomotion modulation on average. The parvalbumin-positive population of inhibitory RSC neurons was particularly strongly modulated by locomotion, both in the dark and during visual stimulation, and this modulation was strikingly stably maintained over the interval of weeks examined. This could provide a mechanism of distinct modes of processing in RSC dependent upon behavioural context, whereby functionally distinct subpopulations are either facilitated or supressed depending upon locomotion state.

Given the diverse roles ascribed to RSC, the similarities to primary visual cortex with respect to the organisation of visually response neurons and their response properties is surprising. In common with V1, many RSC neurons respond to particular positions, orientations and directions of flow of visual stimuli, are topographically organised into a retinotopic map, and exhibit stable visual preferences over periods of weeks. It will be of interest to determine how other modalities of sensory information are functionally represented and topographically organised, and how these maps are integrated with those of visually responsive cells.

In summary, our results show that a subpopulation of RSC neurons surprisingly stably relay retinotopic, orientation and direction information about visual cues as well as locomotion signals. It will be important in future to determine how these view-dependent visual cues representations interact with and contribute to view-independent or allocentric RSC representations. One emergent issue relates to evidence that the RSC may be involved in the identification of stable landmarks in an environment (Auger et al. 2012, 2015; Auger and Maguire 2018; Mitchell et al. 2018). While the separation between view-dependent and view-invariant representations enables different forms of scene and object-based information (Wood 2009), it is their co-operation that would help the formation of landmarks to assist flexible navigation.

## Author contributions

Experiments were performed by AP. Surgical procedures were developed and performed by AP, AV and AR. Experiments were designed by AP, JA, FS, SV, AN and AR. Funding was acquired by JA, FS, SV, AN and AR. Data analysis and visualisation was performed by AR and WMC. Original draft of the manuscript was written by AR. All authors contributed to the review and editing of the manuscript.

## Acknowledgments

This work was supported by a BBSRC research grant awarded to JA, AN, SV and FS (BB/L021005/1) and a Sêr Cymru fellowship (80762-CU-080) to AR. For the use of GCaMP6S we acknowledge Vivek Jayaraman, Rex A. Kerr, Douglas S. Kim, Loren L. Looger, Karel Svoboda from the GENIE Project, Janelia Farm Research Campus, Howard Hughes Medical Institute.

## Supplementary Material

**Figure S1.**
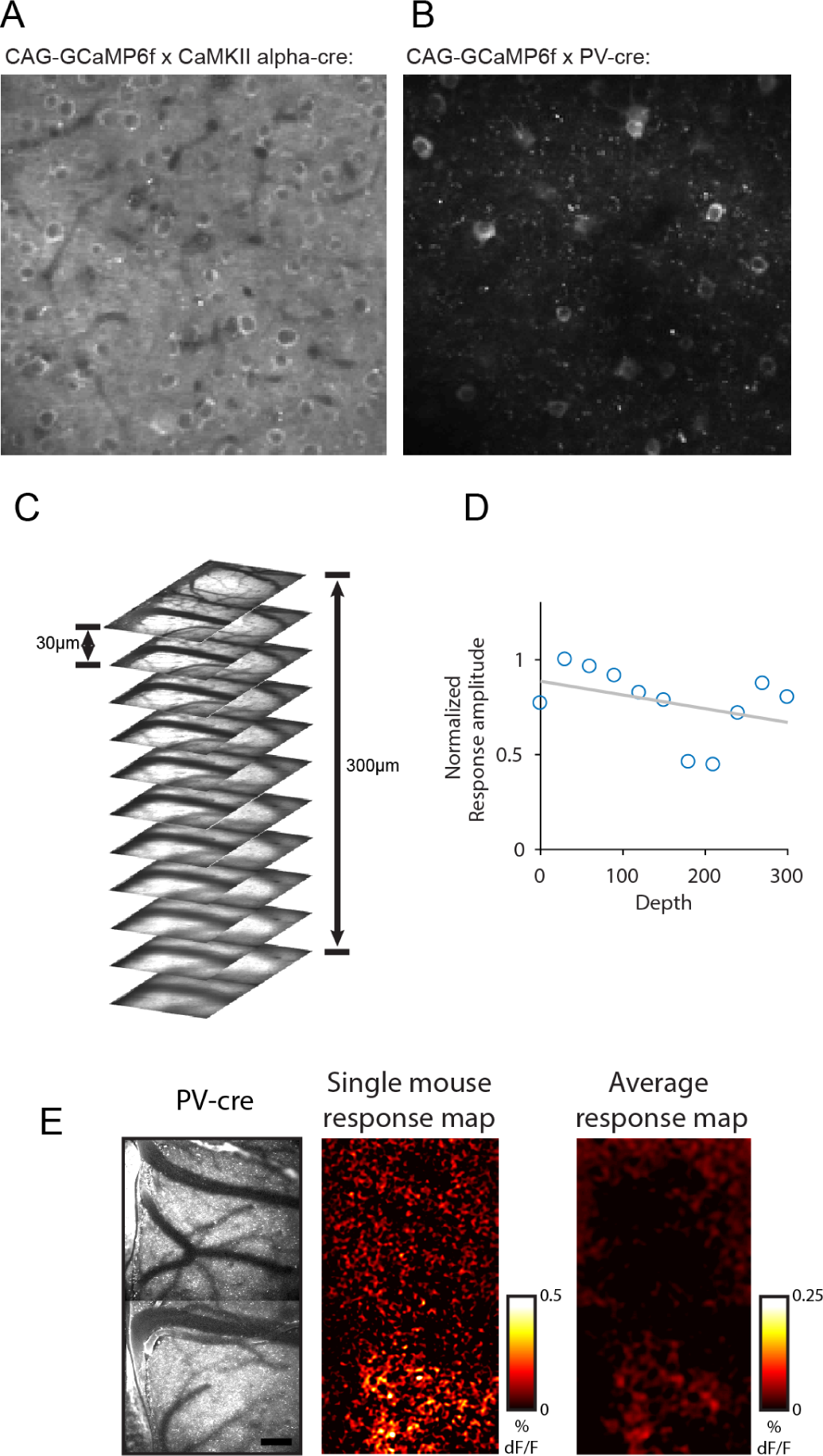
Associated with figure 1. (A-B) Typical field of view of GCaMP6f labelled neurons obtained from cross of CAG-GCaMP6f x CaMKII-cre mice (A) and from cross of CAG-GCaMP6f x PV-cre mice (B). (C) Typical range of depths collected from each animal during wide field recordings. (D) Normalized response amplitude as a function of depth from surface of brain. (E) Example visual response map from a mouse expressing GCaMP6f in PV neurons and mean response map averaged across 5 mice. Scale bar 160µm.

**Figure S2.**
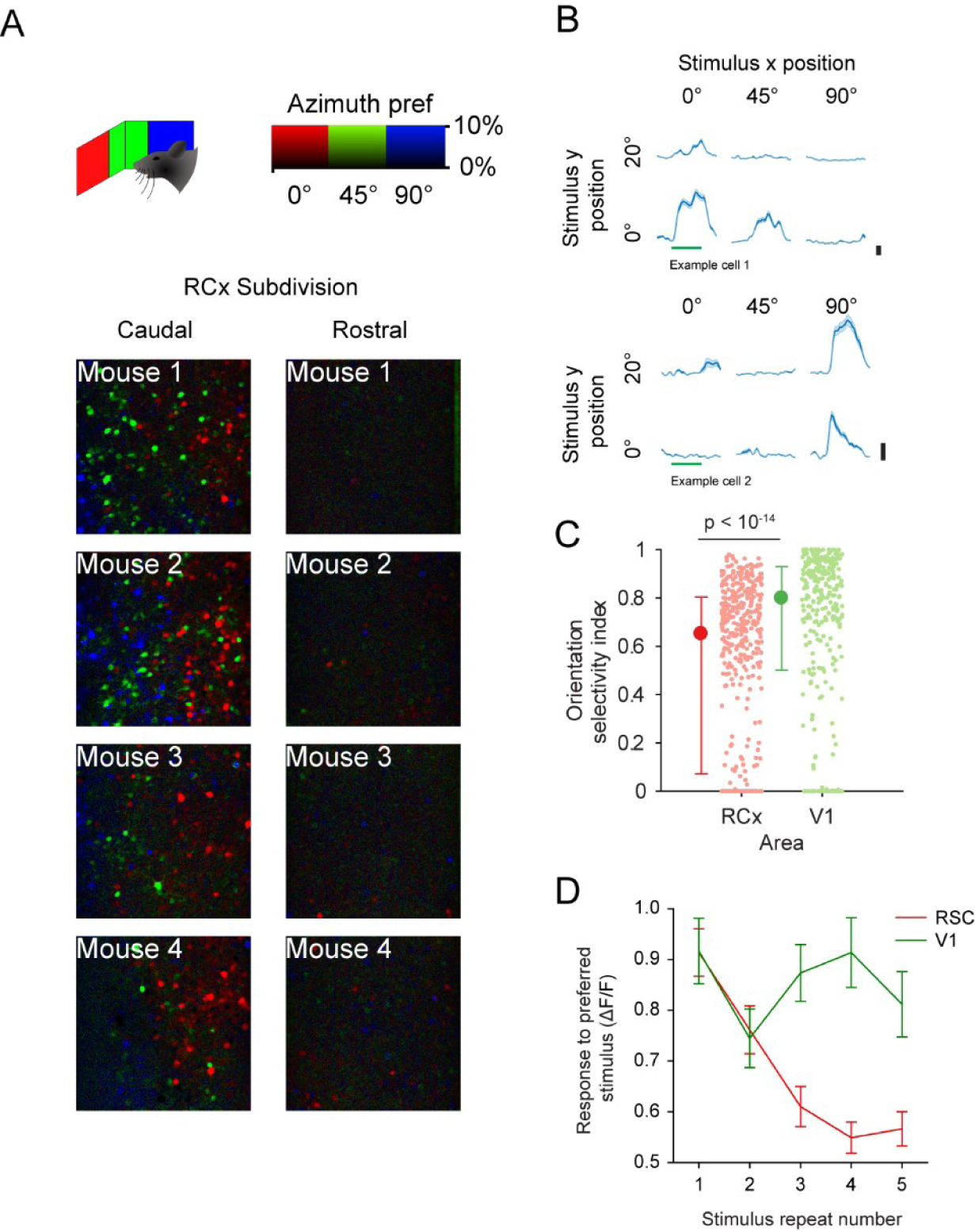
Additional figures associated with figure 2. (A) Pixel-wise response maps from 4 mice showing visual responses in caudal (left) but not rostral (right) retrosplenial cortex. (B) Example mean responses of two neurons with distinct spatial tuning, recorded from the same field of view. (C) Distribution of orientation selectivity indices from all recorded CaMKII cRSC neurons. (D) Rate of adaptation of response to preferred stimulus of RSC and V1 neurons.

**Figure S3.**
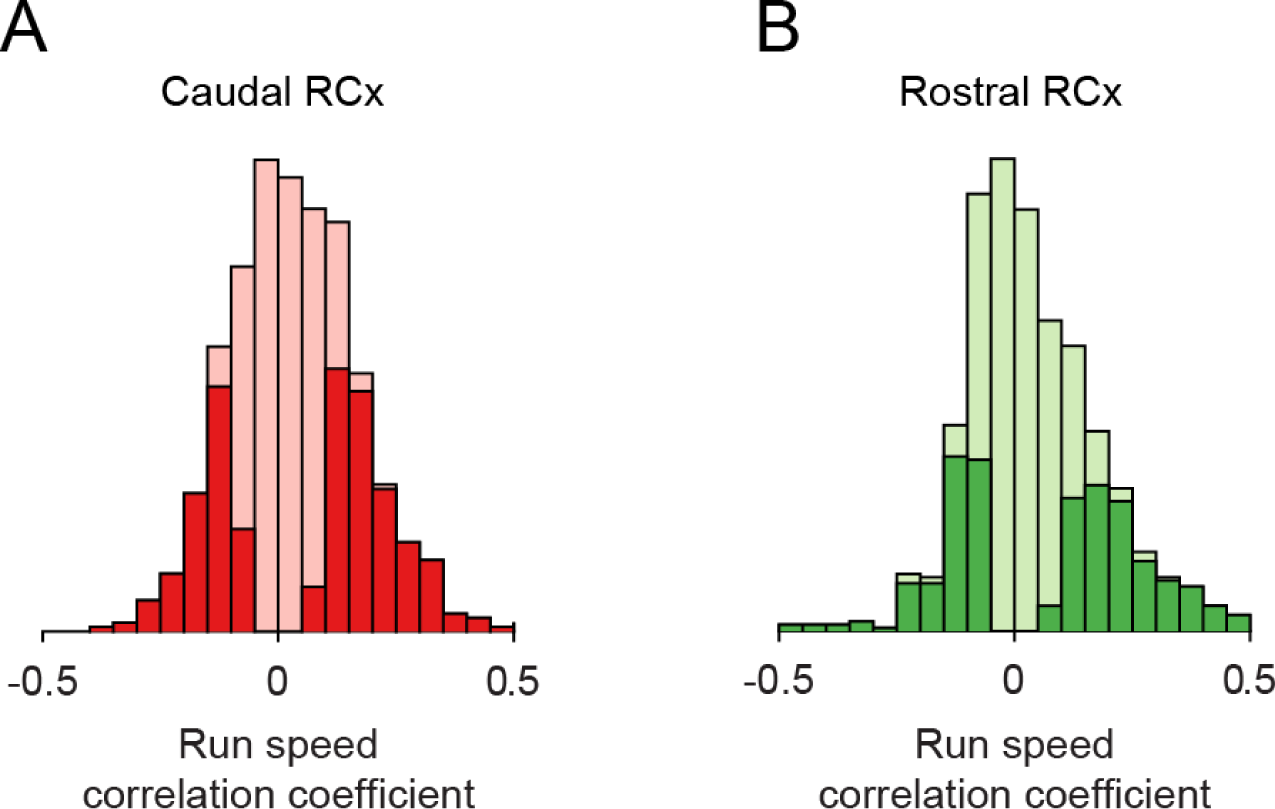
Associated with figure 3. (A-B) The distribution of run speed correlation coefficients of CaMKII neurons in cRSC and rRSC is similar.

**Figure S4.**
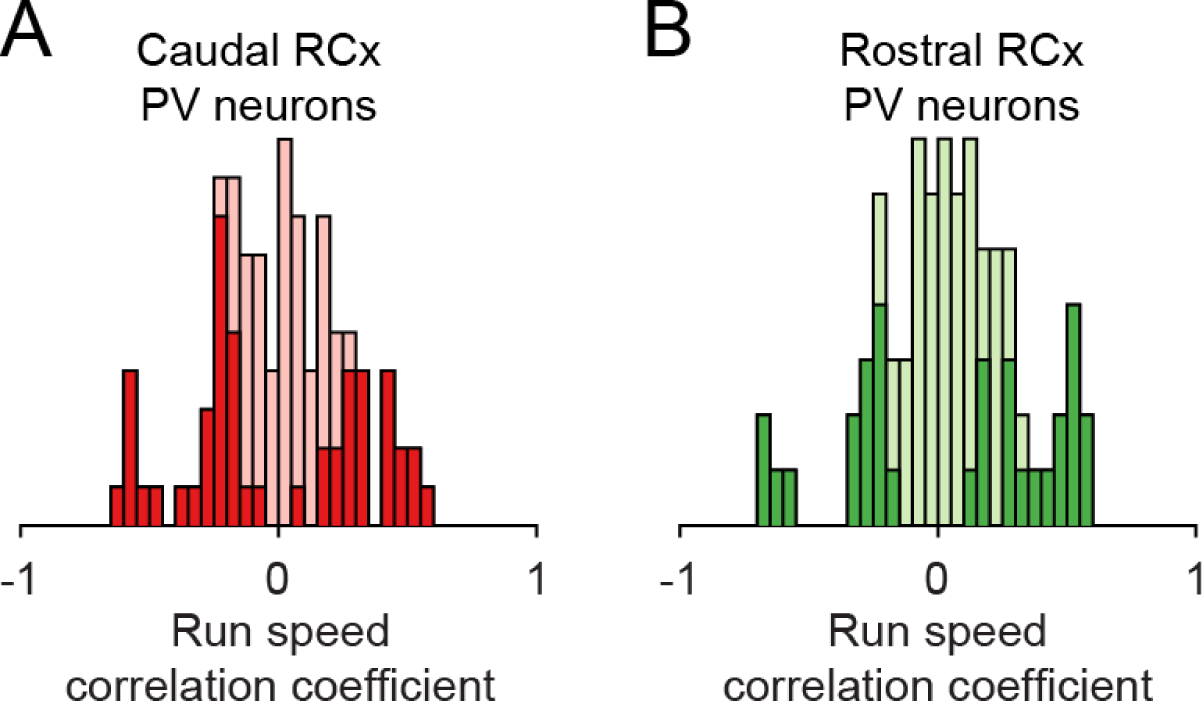
Associated with figure 4. (A-B) The distribution of run speed correlation coefficients of PV neurons in cRSC and rRSC is similar.

**Figure S5.**
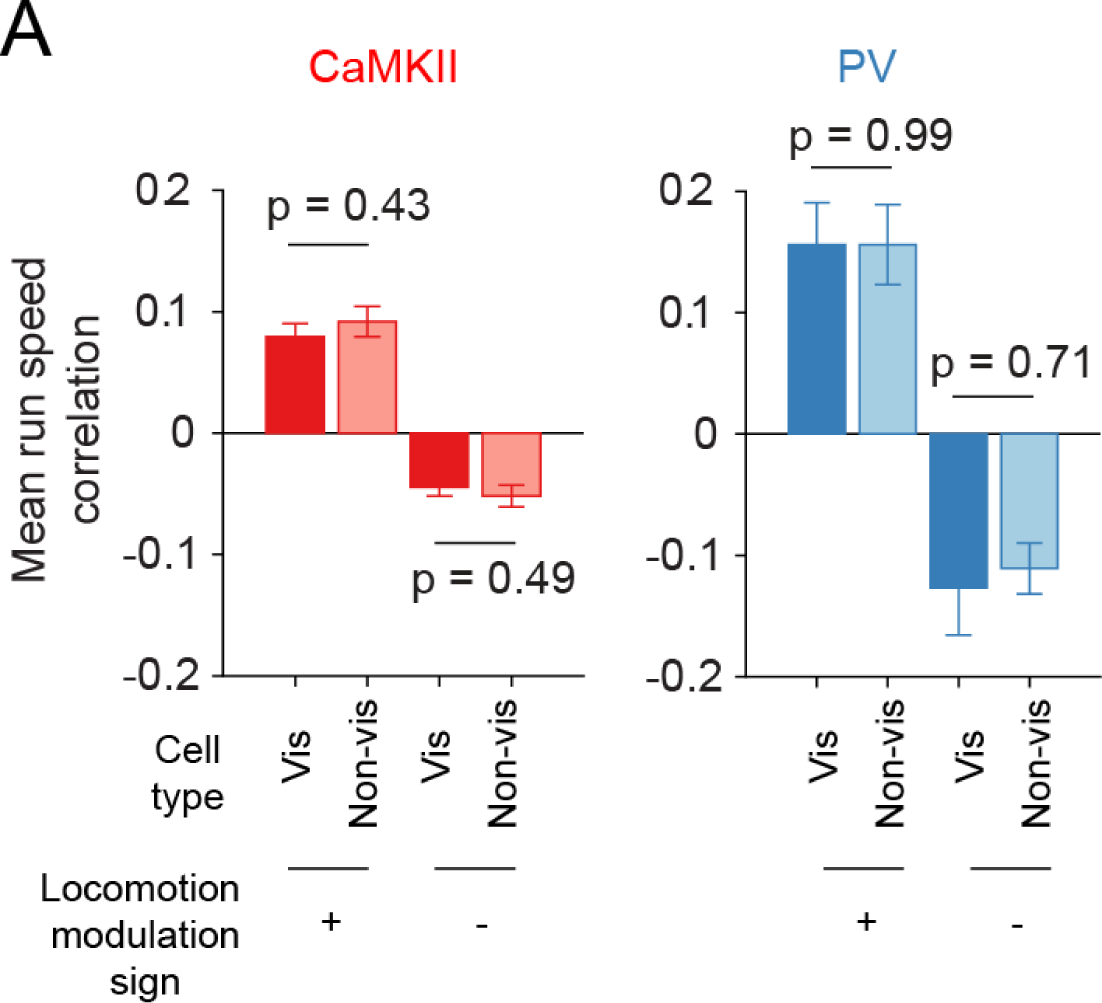
Associated with figure 5. (A) Visually responsive and non-visually responsive neurons have similar average run speed correlation coefficients.

**Figure S6.**
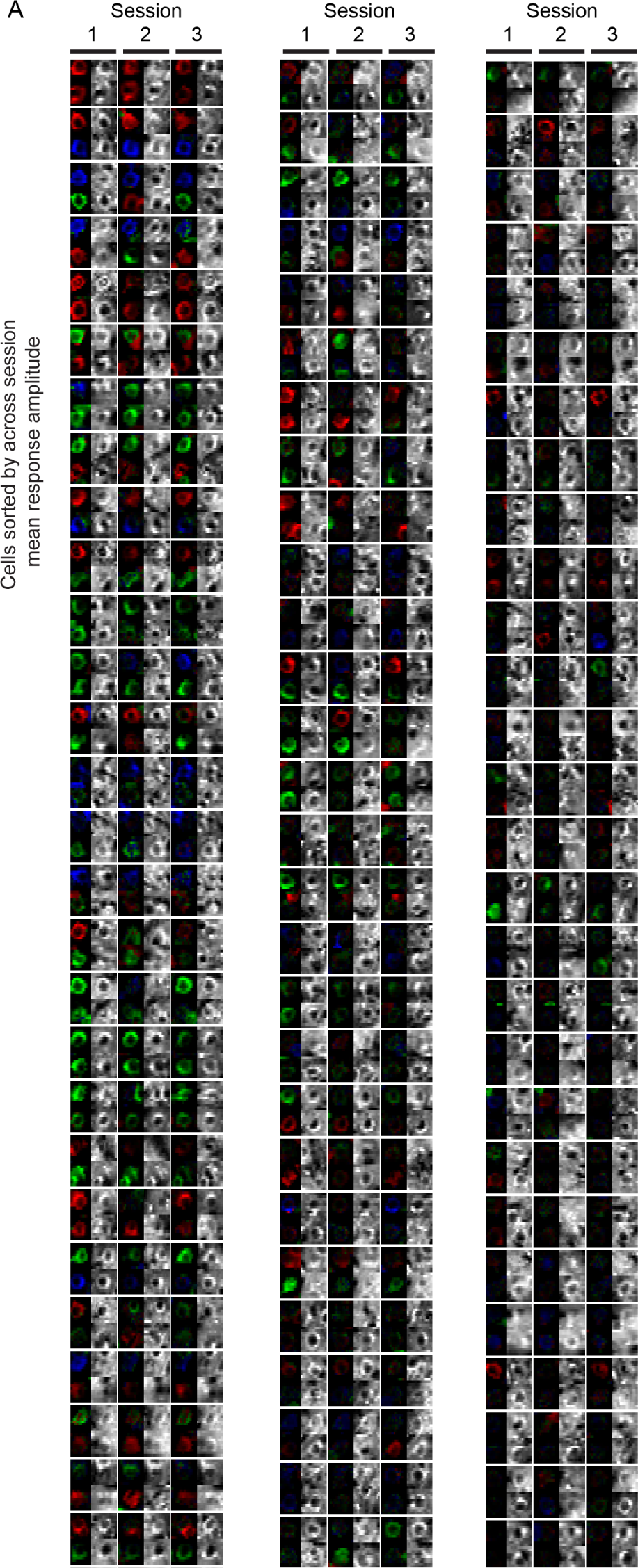
Associated with figure 6. All manually verified longitudinally imaged cRSC CaMKII neurons, colour coded with azimuth stimulus positional preference.

**Figure S7.**
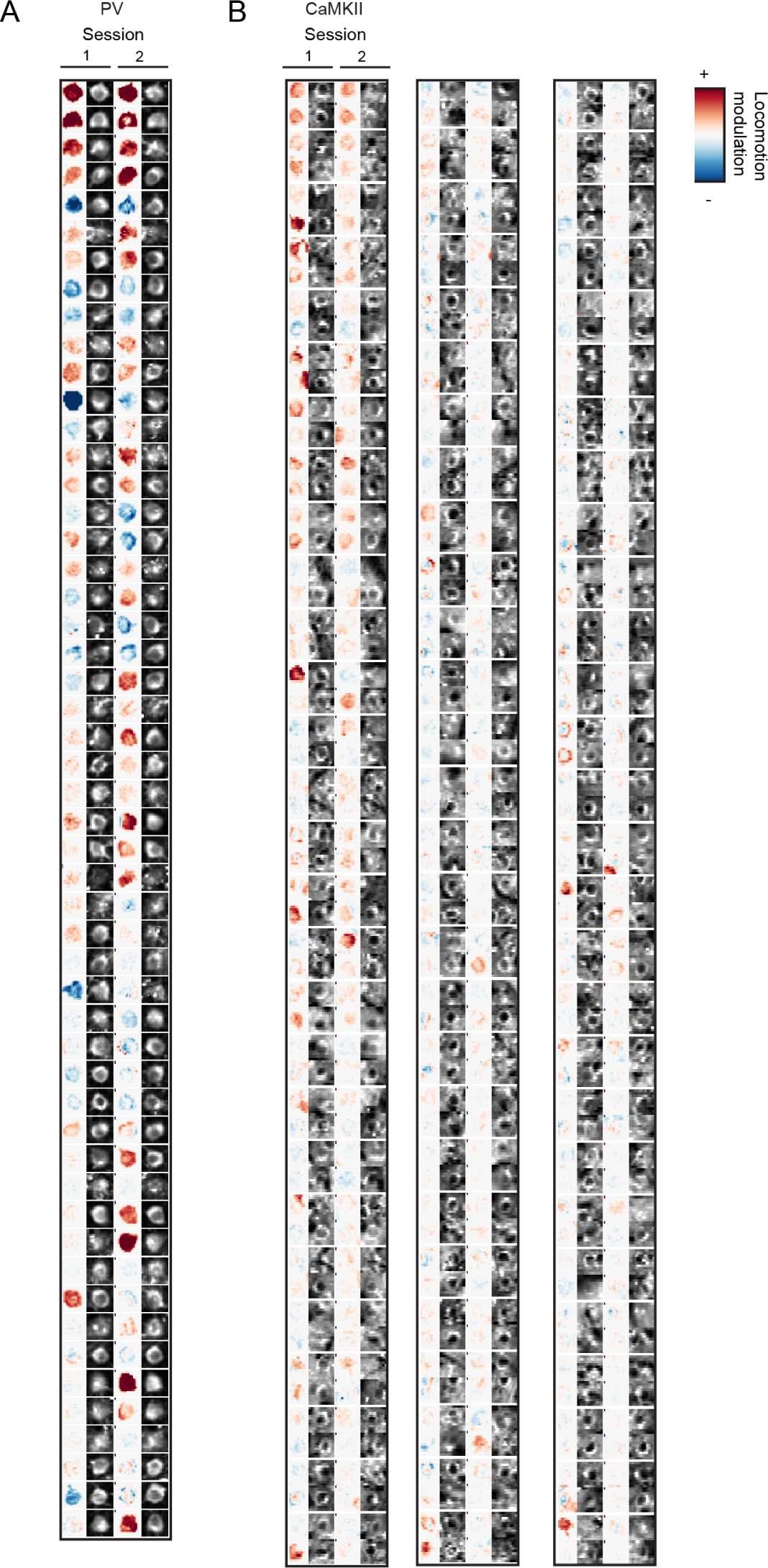
Associated with figure 7. (A) All manually verified longitudinally imaged cRSC PV neurons colour coded with locomotion modulation, where red indicates positive modulation and blue negative. (B) As in (A), but for cRSC CaMKII neurons

